# The impact of leadered and leaderless gene structures on translation efficiency, transcript stability, and predicted transcription rates in *Mycobacterium smegmatis*

**DOI:** 10.1101/857003

**Authors:** Tien G. Nguyen, Diego A. Vargas-Blanco, Louis A. Roberts, Scarlet S. Shell

**Author notes:** These authors contributed equally (Tien G. Nguyen, Diego A. Vargas-Blanco). Address correspondence to Scarlet S. Shell,.

## Abstract

Regulation of gene expression is critical for the pathogen *Mycobacterium tuberculosis* to tolerate stressors encountered during infection, and for non-pathogenic mycobacteria such as *Mycobacterium smegmatis* to survive stressors encountered in the environment. Unlike better studied models, mycobacteria express ∼14% of their genes as leaderless transcripts. However, the impacts of leaderless transcript structures on mRNA half-life and translation efficiency in mycobacteria have not been directly tested. For leadered transcripts, the contributions of 5’ UTRs to mRNA half-life and translation efficiency are similarly unknown. In both *M. tuberculosis* and *M. smegmatis*, the essential sigma factor, SigA, is encoded by an unstable transcript with a relatively short half-life. We hypothesized that *sigA*’s long 5’ UTR caused this instability. To test this, we constructed fluorescence reporters and then measured protein abundance, mRNA abundance, and mRNA half-life. From these data we also calculated relative transcription rates. We found that the *sigA* 5’ UTR confers an increased transcription rate, a shorter mRNA half-life, and a decreased translation rate compared to a synthetic 5’ UTR commonly used in mycobacterial expression plasmids. Leaderless transcripts produced less protein compared to any of the leadered transcripts. However, translation rates were similar to those of transcripts with the *sigA* 5’ UTR, and the protein levels were instead explained by lower transcript abundance. A global comparison of *M. tuberculosis* mRNA and protein abundances failed to reveal systematic differences in protein:mRNA ratios for natural leadered and leaderless transcripts, consistent with the idea that variability in translation efficiency among mycobacterial genes is largely driven by factors other than leader status. The variability in mRNA half-life and predicted transcription rate among our constructs could not be explained by their different translation efficiencies, indicating that other factors are responsible for these properties and highlighting the myriad and complex roles played by 5’ UTRs and other sequences downstream of transcription start sites.

## INTRODUCTION

The pathogen *Mycobacterium tuberculosis* has evolved numerous strategies to survive in different niches within the human host. Bacterial adaptation to these harsh environments is usually achieved by gene regulation, both transcriptionally and post-transcriptionally. While promoters play critical roles in gene regulation, other gene features and mechanisms have additional important regulatory roles. One such important gene feature is the 5’ untranslated region (5’ UTR), which contains the Shine-Dalgarno (SD) sequence within the ribosome binding site (RBS), and therefore can serve as a translation regulator [1-5]. For example, 5’ UTR interactions with *cis* and *trans* elements, such as complementary sequences within the UTR or coding sequence, small RNAs (sRNAs), and RNA-binding proteins, can modulate protein synthesis by blocking or improving accessibility to the RBS [6-9]. Importantly, it has been shown in *Escherichia coli* and other bacteria that transcription and translation are physically coupled, and thus 5’ UTR-mediated modulation of translation could have repercussions on transcription rate as well [10-14]. Translation blocks in *M. smegmatis* have been shown to decrease transcription as well [15], suggesting that transcription-translation coupling occurs in mycobacteria, although the extent and consequences are unknown. 5’ UTRs can also regulate gene expression by altering mRNA turnover rates. This can be a consequence of altered translation rates, as impairments to translation often lead to faster mRNA decay [16-22]. In other cases, mRNA stability is directly affected by sRNA binding to 5’ UTRs or by UTR secondary structure [9, 23-28]. In *E. coli*, the half-life of the short-lived transcript *bla* can be significantly increased when its native 5’ UTR is replaced with the 5’ UTR of *ompA*, a long-lived transcript [29-31]. Conversely, deletion of *ompA*’s native 5’ UTR decreased its half-life by 5-fold [30]. The longevity conferred by the *ompA* 5’ UTR was attributed to the presence of a non-specific stem-loop as well as the specific RBS sequence [30-32]. Secondary structure formation in 5’ UTRs has been shown to play a major role in transcript stability in other bacteria as well, such as for *ermC* in *Bacillus subtilis* [33, 34] and *pufBA* in *Rhodobacter capsulatus* [35-37]. Moreover, obstacles that hinder the linear 5’ scanning function of RNase E (a major RNase in *E. coli* and mycobacteria) can prevent access to downstream cleavage sites, increasing transcript half-life [38]. Such obstacles include the 30S ribosomal subunit bound to an SD-like site far upstream of the translation start site in one case [39]. UTRs can also contain binding sites for the global regulator CsrA, which can both promote and prevent mRNA decay in *E. coli* [40]. Although effects of 5’ UTRs on mRNA stability, translation, and transcription rate have been widely studied in more common bacterial systems, there is a paucity of information of the regulatory effects of 5’ UTRs in mycobacteria.

Compared to *E. coli* and most other well-studied bacteria, mycobacteria possess a large number of leaderless transcripts; approximately 14% of annotated genes are leaderless in both *M. smegmatis* and *M. tuberculosis* [41-43]. Studies in *E. coli* have shown that translation of leadered and leaderless transcripts is functionally distinct [44-50], suggesting fundamental differences in their mechanisms of regulation. In contrast to *E. coli*, where leaderless transcripts are generally translated less efficiently [42, 51-53], leaderless transcripts in mycobacteria appear to be translated robustly [42, 43]. However, direct comparisons of translation rates for leadered vs leaderless transcripts in mycobacteria have yet to be reported.

Among leadered transcripts, 5’ UTR lengths vary. We hypothesized that longer 5’ UTRs were more likely to play regulatory roles through modulation of translation, transcription rate, and mRNA turnover. One such long-leadered transcript in both *M. tuberculosis* and *M. smegmatis* encodes sigma factor alpha (*sigA*), the primary sigma factor in mycobacteria [54, 55]. Here we used the mycobacterial model *M. smegmatis* and a series of yellow fluorescent protein (YFP) reporters to investigate the effects of the *sigA* 5’ UTR as well as leaderless gene structures on transcription, translation, and mRNA half-life. We found that the *sigA* 5’ UTR caused lower translation efficiency, reduced mRNA half-life, and a higher predicted transcription rate compared to a control 5’ UTR. Leaderless transcripts were translated at similar rates as transcripts bearing the *sigA* 5’ UTR and had similar half-lives, but appeared to be transcribed less efficiently, leading to lower steady-state mRNA and protein abundances. Our results highlight the potential of 5’ UTRs to affect transcription efficiency as well as translation and mRNA half-life, and support the idea that leaderless translation can be either more or less efficient than leadered translation in mycobacteria, depending on the characteristics of the leader.

## RESULTS

### Validation of the *sigA* 5’ UTR boundaries

Transcription start site mapping has defined the 5’ ends of 5’ UTRs on a transcriptome-wide basis in both *M. smegmatis* and *M. tuberculosis* [41, 42]. Using annotated translation start sites to define the 3’ ends of the 5’ UTRs, the median 5’ UTR lengths in *M. smegmatis* and *M. tuberculosis* are 64 and 71 nt, respectively (Fig. 1A and table S1, [41, 42]). The 5’ UTR length distributions are skewed, with a mode of approximately 40 nt (Fig. 1A). We hypothesized that longer-than-average 5’ UTRs are more likely to have regulatory roles, and sought to investigate the role of the 5’ UTR of the *M. smegmatis sigA* gene. The *M. tuberculosis sigA* 5’ UTR is also predicted to be longer than the median (128 nt). To ensure that the predicted 5’ UTR boundaries of *M. smegmatis sigA* were correct, we experimentally validated the predicted start codon at NC_008596 genome coordinate 2827625, which resulted in a 123 nt UTR. A second GTG codon 39 nt downstream at 2827625 also had an appropriately positioned Shine-Dalgarno-like sequence and could conceivably be used as a start codon. We therefore made reporter constructs in which the strong constitutive promoter p_myc1_*tetO* [56] drove expression of a transcript containing the *sigA* 5’ UTR and the sequence encoding YFP, with a C-terminal 6xHis tag and an N-terminal fusion of the sequence encoded by the first 54 nt of the annotated *sigA* coding sequence. We then individually mutated each of two putative GTG start codons to GT<u>C</u> (Fig. 1B). Mutations of the first GTG to GT<u>C</u> reduced fluorescence to levels indistinguishable from auto-fluorescence in a strain that lacked the *yfp* gene (Fig. 1C and D). In contrast, mutation of the second GTG to GTC reduced fluorescence to an intermediate level (Fig. 1C and D). We therefore concluded that the first GTG is likely to be the predominant site of translation initiation, while the second GTG may affect expression levels but is not by itself sufficient to produce above-background expression. For subsequent experiments, we considered the first GTG to be the most likely start of the coding sequence, and thereby define the *sigA* 5’ UTR as 123 nt in length.

**Figure 1.**
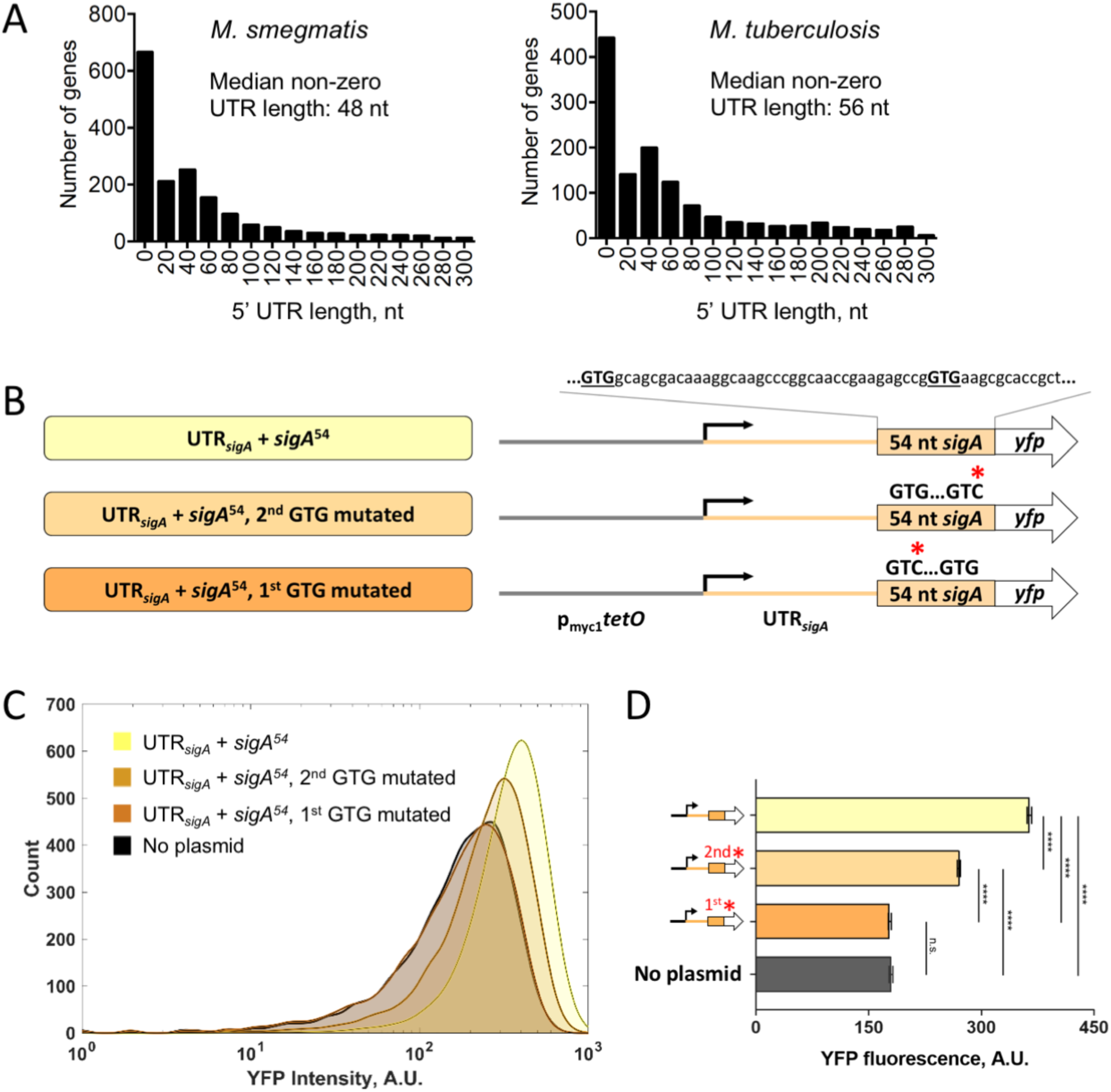
The *M. smegmatis sigA* gene has a longer-than-typical 5’ UTR. **A:** Distributions of 5’ UTR lengths for *M. smegmatis* and *M. tuberculosis* genes reported to be transcribed from a single TSS [41, 42]. **B:** Constructs to confirm the predicted *sigA* translation start site. p_myc1_*tetO* was described in [56]. UTR_sigA_ denotes the 123 nt sequence between the experimentally determined TSS [41] and the annotated translation start site. **C:** Flow cytometry with YFP-expressing constructs diagrammed in B. **D:** Median fluorescence intensities determined by flow cytometry. Error bars denote 95% CI. Fluorescence intensities were compared by Kruskal-Wallis test followed by Dunn’s multiple comparisons test. ****, *p* < 0.0001; ns, *p* > 0.05.

### The initial portion of the *sigA* coding sequence affects mRNA half-life and predicted transcription rate

To capture 5’ UTR-dependent effects on transcription, mRNA stability, and translation, we sought to investigate the role of the *sigA* 5’ UTR (UTR_*sigA*_) in the context of a *yfp* transcript. UTR-mediated regulation of translation sometimes involves base pairing of 5’ UTR sequences with elements in the early portion of the coding sequence. Thus, we decided to include in our investigation the first 54 nt from the coding region of *sigA* (*sigA*^54^). To determine if *sigA*^54^ alone affected expression, we compared fluorescence from our YFP reporters with or without the *sigA*^54^ N-terminal tag, independent from UTR_*sigA*_. Transcription was driven by the p_myc1_*tetO* promoter for these and all constructs used in this study. While this semi-synthetic promoter contains TetR binding sites, the strains used in this study did not encode the corresponding Tet repressor and the promoter was therefore constitutively active. Where indicated, constructs included the p_myc1_*tetO* associated 5’ UTR (UTR_pmyc1*tetO*_) as initially described by [56]. To ensure that expression initiated only from the annotated promoter and not from spurious promoter-like sequences in UTR_pmyc1*tetO*_ or *sigA*^54^, we built a control strain in which nt −53 through −1 of the promoter were deleted (Δp_myc1_*tetO*) (Fig. 2A). As shown in Fig. 2B, background fluorescence intensity was indistinguishable between Δp_myc1_*tetO* and a strain lacking the YFP cassette completely.

**Figure 2.**
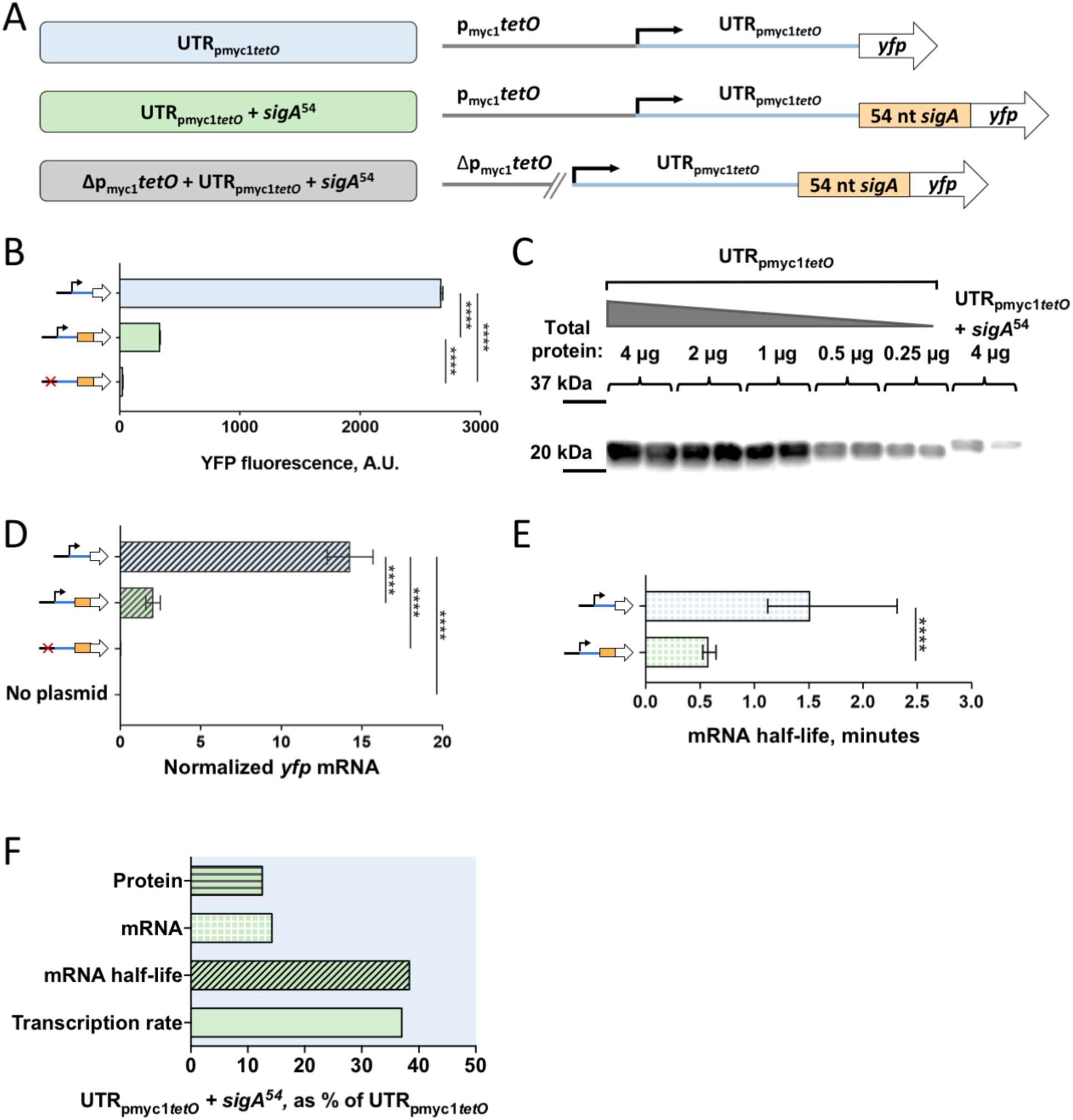
The first 54 nt of the *sigA* coding sequence affects transcription rate and mRNA half-life. **A:** Constructs transformed into *M. smegmatis* to determine the impact of the first 54 nt of the sigA coding sequence (*sigA*^*54*^) on expression of a YFP reporter. **B:** Median YFP fluorescence of strains bearing the constructs in A, determined by flow cytometry. Error bars denote 95% CI. Strains were compared by Kruskal-Wallis test followed by Dunn’s multiple comparisons test. ****, *p* < 0.0001. **C:** Lysates from strains bearing constructs with and without *sigA*^*54*^ were subject to western blotting to detect the C-terminal 6x His tag on the YFP. The mass of total protein loaded per lane is stated. **D:** *yfp* mRNA abundance for strains bearing the indicated constructs, determined by qPCR and normalized to expression of endogenous *sigA*. Error bars denote SD. Strains were compared by ANOVA with Tukey’s HSD. **E:** The half-lives of *yfp* mRNA produced from the indicated constructs were measured. Error bars denote 95% CI. Half-lives were compared using linear regression analysis (*n*=3). **F:** Protein abundance, mRNA abundance, mRNA half-life, and calculated transcription rate for the construct containing *sigA*^*54*^ are shown as a percentage of the values produced by a construct that lacks *sigA*^*54*^, but is otherwise identical.

We first tested the impact of *sigA*^54^ on YFP fluorescence intensity using UTR_pmyc1*tetO*._ Interestingly, the *sigA*^54^ strain was ∼nine-fold less fluorescent than the strain in which YFP lacked this N-terminal tag (Fig. 2B). To confirm that the reduced YFP fluorescence in the presence of *sigA*^54^ indeed reflected reduced protein levels rather than altered YFP structure or intrinsic fluorescence, we measured protein levels directly by western blotting. The western blotting data were consistent with the flow cytometry result, showing an approximately 16-fold reduction of YFP levels with the inclusion of *sigA*^54^ compared to the no-*sigA*^54^ strain (Fig. 2C).

To assess if the presence of *sigA*^54^ affected *yfp* transcript levels, we conducted quantitative PCR (qPCR) for the same set of strains. Indeed, *sigA*^54^*yfp* levels were approximately six-fold lower than those of the *yfp* strain (Fig. 2D). This suggested that the decrease in YFP protein levels was due primarily to a reduction in *yfp* mRNA.

We then wondered if *sigA*^54^ affected transcript abundance by increasing the rate of transcript decay or by decreasing the rate of transcription. Thus, we determined mRNA half-life for *yfp* with and without *sigA*^54^. As shown in Fig. 2E, we estimated the half-life of *yfp* alone to be ∼1.5 min, and the half-life of *yfp* N-tagged with *sigA*^54^ to be ∼0.6 min. We concluded that the first 54 nt of *sigA* made the *yfp* transcript more susceptible to degradation. Knowing the abundance and decay rate of a transcript, the rate of transcription can be predicted mathematically [57]. The insertion of *sigA*^54^ as an N-terminal tag for YFP appeared to reduce the *yfp* transcription rate by approximately 55%, suggesting that the reduced steady-state transcript abundance resulted from a combination of slower transcription and faster decay (Fig. 2F).

### The *sigA* 5’ UTR affects transcript half-life, translation, and predicted transcription rate

In order to assess the effects of UTR_*sigA*_ on transcription, mRNA stability, and translation, we replaced UTR_pmyc1*tetO*_ with UTR_*sigA*_ in our *sigA*^54^*yfp* reporters as shown in Fig. 3A. The presence of UTR_*sigA*_ led to an approximately two-fold reduction in YFP fluorescence intensity when compared to the UTR_pmyc1*tetO*_ reporter strain (Fig. 3B). We wondered if the reduction in fluorescence attributed to UTR_*sigA*_ was caused by reduced *yfp* transcript abundance. However, qPCR revealed equivalent transcript levels for strains with UTR_*sigA*_ and UTR_pmyc1*tetO*_ (Fig 3C), indicating that the reduced protein levels were more likely a consequence of reduced translation efficiency. Interestingly, *yfp* mRNA half-life was reduced to 0.28 min by the presence of UTR_*sigA*_ (Fig. 3D), indicating that a higher transcription rate is required to maintain the steady-state mRNA abundance that we observed (Fig. 3E). Taken together, our findings suggest that UTR_*sigA*_ affects transcription, transcript decay, and translation. In Fig. 3E we summarize these results as percentages of *yfp* transcription rate, mRNA abundance, mRNA half-life, and YFP protein levels relative to the UTR_pmyc1*tetO*_ *sigA*^54^ strain.

**Figure 3.**
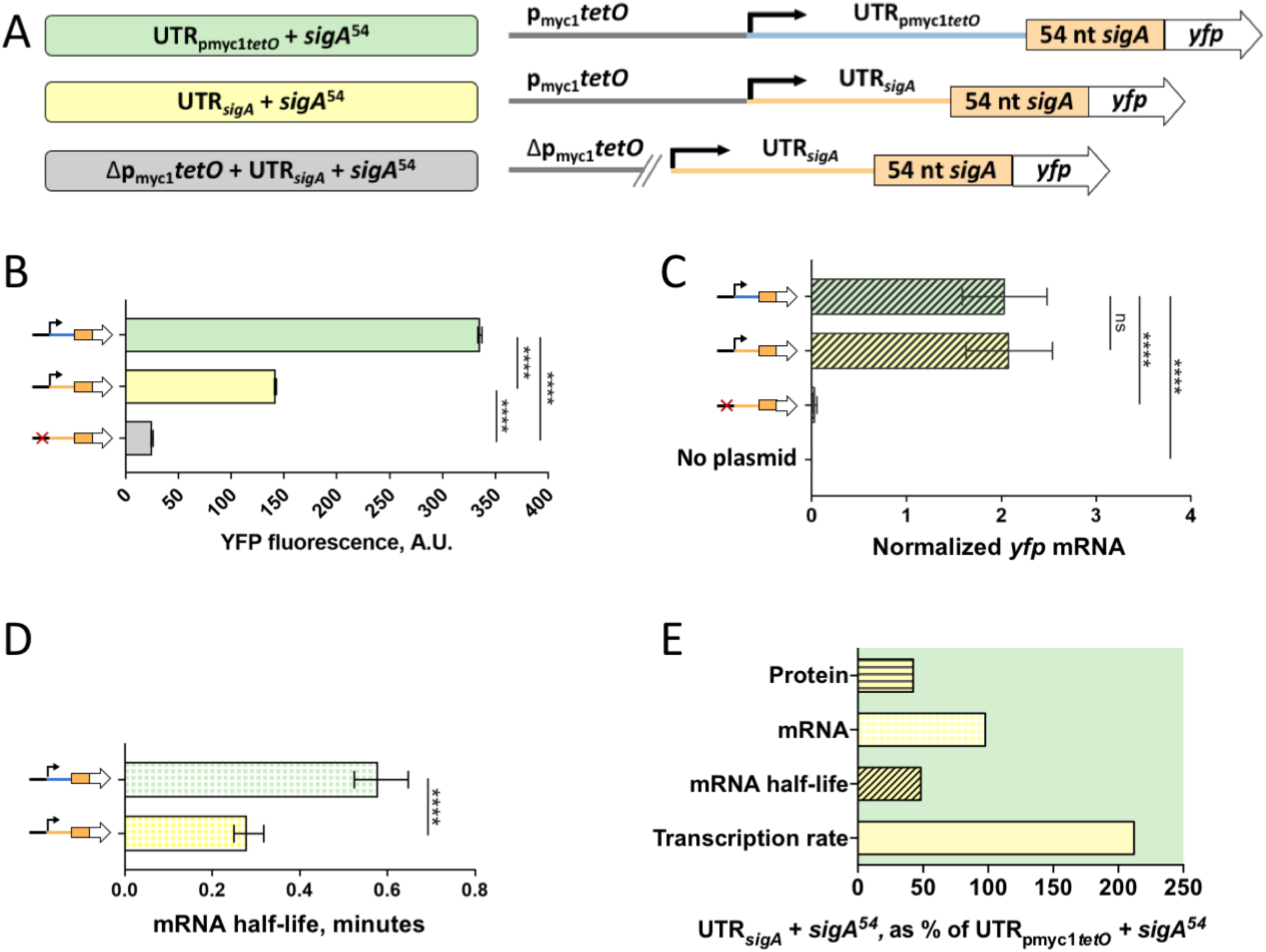
The *sigA* 5’ UTR affects translation efficiency, mRNA half-life, and transcription rate. **A:** Constructs transformed into *M. smegmatis* to determine the impact of the *sigA* 5’ UTR on expression of a YFP reporter. **B:** Median YFP fluorescence of strains bearing the constructs in A, determined by flow cytometry. Error bars denote 95% CI. Strains were compared by Kruskal-Wallis test followed by Dunn’s multiple comparisons test. ****, *p* < 0.0001. **C:** *yfp* mRNA abundance for strains bearing the indicated constructs, determined by qPCR and normalized to expression of endogenous *sigA*. Error bars denote SD. Strains were compared by ANOVA with Tukey’s HSD.. ****, p < 0.0001; ns, *p* > 0.05. **D:** The half-lives of *yfp* mRNA produced from the indicated constructs were measured. Error bars denote 95% CI. Half-lives were compared using linear regression analysis (*n*=3). **E:** Protein abundance, mRNA abundance, mRNA half-life, and calculated transcription rate for the construct containing the *sigA* 5’ UTR are shown as a percentage of the values produced by a construct that contains the p_myc1_*tetO*-associated 5’ UTR.

We analyzed the sequences and predicted secondary structures of UTR_*sigA*_ and UTR_pmyc1*tetO*_ to investigate possible causes of the difference in translation efficiency. The ribosome binding sites (RBSs) of these two UTRs have similar degrees of identity to a theoretically perfect mycobacterial Shine-Dalgarno (SD) sequence (the reverse complement of the 3’ end of the 16S rRNA; Fig. S1A). We noted that the spacing between the SD and start codon differed substantially between the two UTRs (Fig. S1A). However, both spacings are common among native *M. smegmatis* transcripts harboring these SD sequences (Fig. S1B), suggesting that the differences in translation efficiency are not due to differences in the favorability of spacing of RBS elements. However, secondary structure predictions by Sfold [58, 59] suggested that the UTR_pmyc1*tetO*_ SD is likely to be in a single-stranded loop while the UTR_*sigA*_ SD is likely to be partially base-paired (Fig. S1C and D), suggesting that differences in SD accessibility could be responsible for the observed differences in translation efficiency.

### Leaderless mRNAs may be transcribed less efficiently

Leaderless transcripts are common in mycobacteria and were found to be associated with reduced protein abundance compared to leadered transcripts with near-consensus Shine-Dalgarno sites [43], suggesting that leaderless translation may be generally less efficient, as was shown in E. coli [51-53]. However, this hypothesis was not experimentally tested in mycobacteria. We therefore built two leaderless *yfp* reporters under the control of the p_myc1_*tetO* promoter, with and without the *sigA*^54^ N-terminal tag (Fig. 4A). When we compared YFP fluorescence between the leadered and leaderless reporters, we found that the leaderless strains were substantially less fluorescent than those containing either 5’ UTR, regardless of the presence of *sigA*^54^ (Fig. 4B). The leaderless constructs also had reduced *yfp* mRNA levels compared to all of the leadered constructs (Fig. 4C). When comparing the leaderless constructs to the UTR_pmyc1*tetO*_ construct, protein levels were decreased to a greater extent than mRNA levels, (Fig. 4D), suggesting that the leaderless mRNAs were indeed translated less efficiently than mRNAs bearing UTR_pmyc1*tetO*_. However, the difference in protein abundance from constructs without leaders and with UTR_*sigA*_ could be explained entirely by the difference in mRNA levels (Fig. 4D), suggesting that leaderless and UTR_*sigA*_-leadered mRNAs are translated with similar efficiencies. These findings indicate that the relative efficiencies of leadered and leaderless translation are dependent on the composition of the leader.

**Figure 4.**
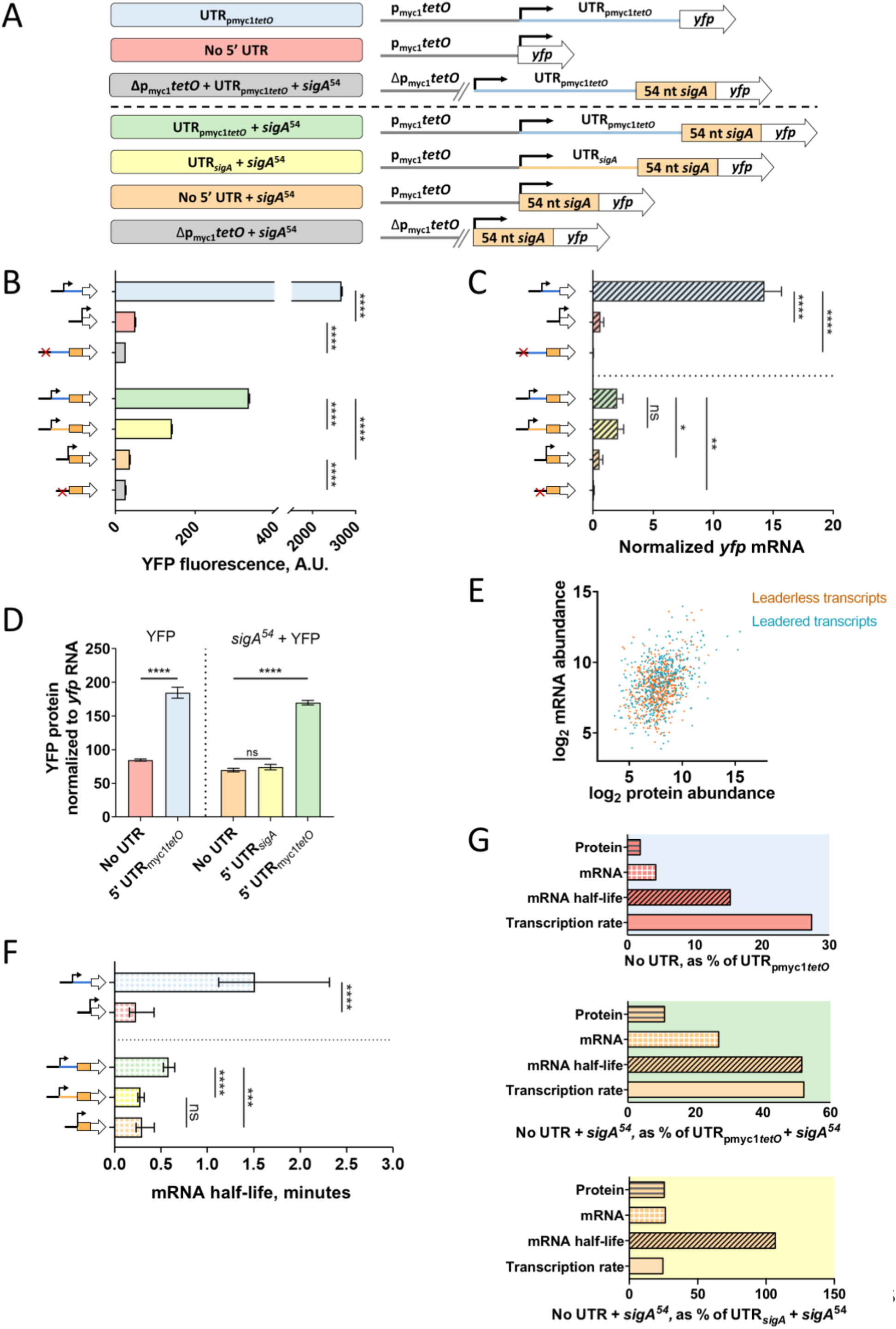
Leaderless transcripts have altered translation efficiencies, mRNA half-lives, and predicted transcription rates compared to leadered controls. **A:** Constructs transformed into *M. smegmatis* to compare leaderless vs. leadered gene structures. **B:** Median YFP fluorescence of strains bearing the constructs in A, determined by flow cytometry. Error bars denote 95% CI. Strains were compared by Kruskal-Wallis test followed by Dunn’s multiple comparisons test. ****, *p* < 0.0001. **C:** *yfp* mRNA abundance for strains bearing the indicated constructs, determined by qPCR and normalized to expression of endogenous *sigA*. Error bars denote SD. Strains were compared by ANOVA with Tukey’s HSD. **D:** Transcripts containing the p_myc1_*tetO*-associated 5’ UTR are translated more efficiently than leaderless transcripts or those containing the *sigA* 5’ UTR. **E:** Published *M. tuberculosis* mRNA abundance [42] and protein abundance [60] levels for genes that have a single TSS and are leaderless or have 5’ UTRs ≥ 15 nt. Protein and mRNA abundance were significantly correlated for both gene structures (*p* < 0.0001, Spearman’s *ρ*). Linear regression analysis revealed that the slopes were statistically indistinguishable (*p* = 0.44). **F:** The half-lives of *yfp* mRNA produced from the indicated constructs were measured. Error bars denote 95% CI. Half-lives were compared using linear regression analysis (*n*=3). **G:** Protein abundance, mRNA abundance, mRNA half-life, and calculated transcription rate for leaderless transcripts compared to transcripts with 5’ UTRs.

To further evaluate the relationship between leader status and translation efficiency, we compared the relative abundances of proteins and mRNAs in *M. tuberculosis* using published quantitative proteomics data [60] and RNAseq data [42]. For both leaderless transcript and transcripts with 5’ UTRs ≥ 15 nt in length, mRNA abundance and protein abundance were significantly correlated (*p* < 0.0001, Spearman’s *ρ*) (Fig. 4E). Linear regression of these correlations revealed that they were statistically indistinguishable for leadered vs leaderless transcripts, consistent with the idea that variability in translation efficiency among mycobacterial genes is largely driven by factors other than presence or absence of a leader.

We wondered if the reduced abundance of the leaderless *yfp* transcripts relative to the UTR_pmyc1*tetO*_-leadered transcripts was associated with reduced mRNA stability. Indeed, *yfp* half-lives for the UTR_pmyc1*tetO*_ leadered transcripts were longer than their leaderless counterparts (Fig. 4F). In contrast, the leaderless transcripts had half-lives similar to the transcript bearing UTR_*sigA*_ (Fig. 4F). Interestingly, leaderless transcripts with and without the *sigA*^54^ N-terminal tag had equivalent half-lives. Taken together, the data indicate that the destabilizing effect of *sigA*^54^ observed in Fig. 2F is dependent on the UTR_pmyc1*tetO*_ present in those constructs.

The predicted *yfp* transcription rates of the leaderless constructs were lower than their leadered counterparts (Fig. 4G). This is consistent with the idea that transcription-translation coupling can cause transcription rates to be altered as a function of translation efficiency [14]. However, the UTR_*sigA*_-leadered transcript appeared to be translated with a similar efficiency as the leaderless constructs (Fig. 4D) and yet had a substantially higher transcription rate. These data suggest that there are additional factors besides promoters and translation efficiency that can affect transcription rate.

### Translation efficiency is a poor predictor of mRNA half-life and transcription rate

The five constructs described above had identical promoters but produced strains that varied widely with respect to protein abundance, mRNA abundance, mRNA half-life, translation efficiency, and predicted transcription efficiency. Given the reported impacts of translation efficiency on mRNA stability in bacteria [16-22], we wondered to what extent the differences in half-life among our constructs were explained by differences in translation efficiency. We defined translation efficiency as:

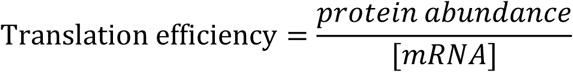

However, the relationship between translation efficiency and mRNA half-life was weak (Fig. 5A), indicating that the variability in mRNA half-life was largely due to other factors. Translation rate has also been reported to affect transcription rate [14], but these two properties did not appear to be correlated in our constructs (Fig. 5B), suggesting that the differences in transcription rate were not a consequence of differences in translation rate.

**Figure 5.**
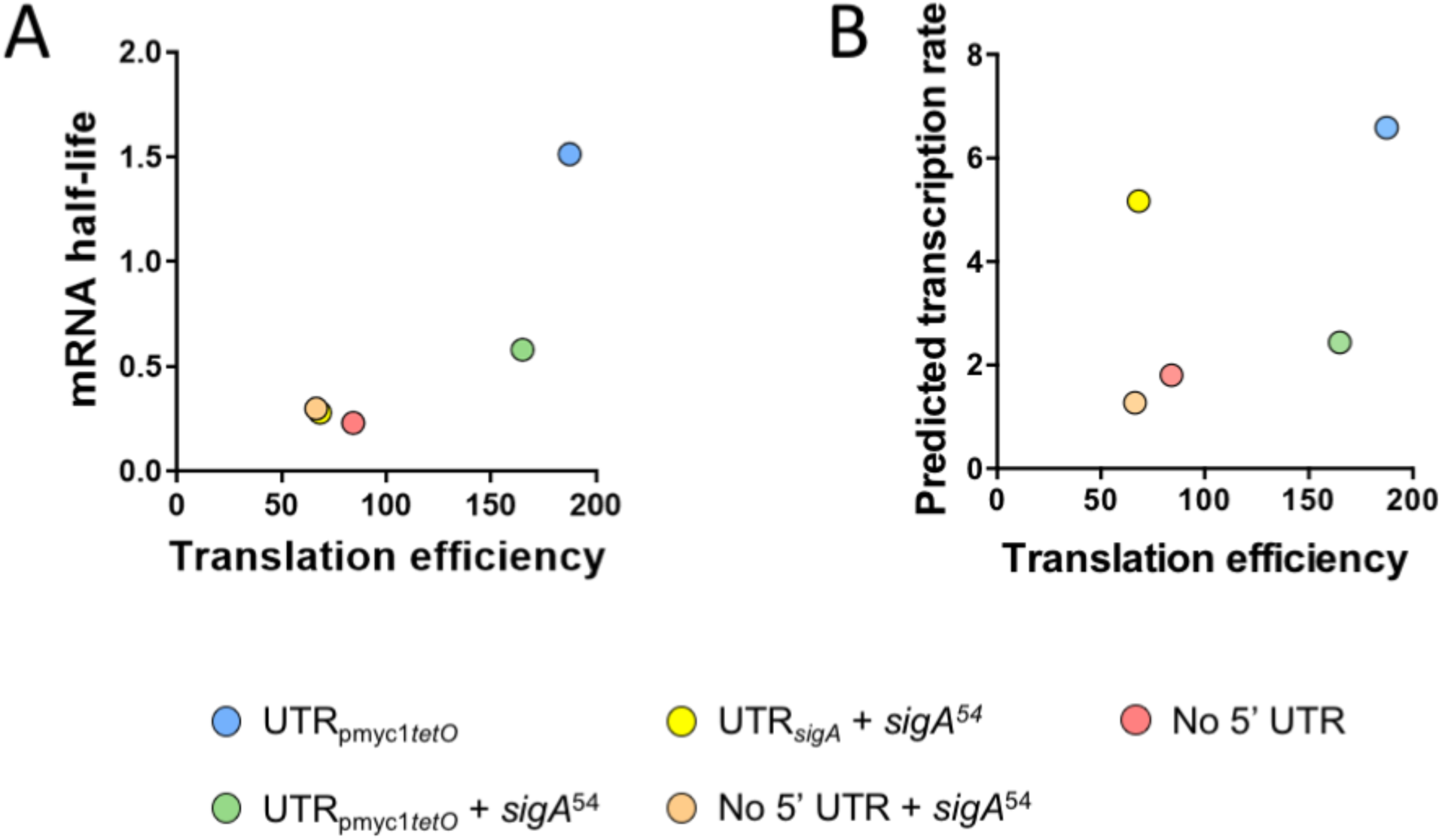
Translation efficiency is poorly correlated with mRNA half-life and predicted transcription rate. Translation efficiency was defined as the ratio of protein abundance to mRNA abundance (arbitrary units). **A:** Variability in mRNA half-life is largely not explained by variability in translation efficiency; and **B:** Variability in predicted transcription rate is uncorrelated with translation efficiency.

## DISCUSSION

*sigA* transcripts in both *M. tuberculosis* and *M. smegmatis* were reported to have relatively short half-lives [15, 61], and we hypothesized that this property was conferred in part by the 5’ UTR. We therefore sought to determine the impacts of the *M. smegmatis sigA* 5’ UTR on expression and mRNA stability. Compared to a 5’ UTR associated with high levels of protein expression and commonly used in mycobacterial expression vectors [56], the *sigA* UTR indeed conferred a shorter half-life as well as reduced translation efficiency. However, the half-life of a *sigA*-leadered transcript was similar to that of a leaderless transcript. Insertion of part of the *sigA* coding sequence as an N-terminal translational fusion to our YFP reporter also caused a reduction in mRNA half-life. These findings suggest the relative instability of the native *sigA* transcript is a product of multiple features, including the 5’ UTR and regions of the coding sequence. However, this effect was not observed for a leaderless version of the translational fusion, indicating the effect is context-dependent.

Our mRNA abundance and half-life data allowed us to calculate predicted transcription rates. It is important to note that these calculated transcription rates reflect the combined contributions of transcription initiation, elongation, and termination, and that our methodology did not allow us to distinguish between these processes. Interestingly, the *sigA* 5’ UTR appeared increase transcription rates when compared to the p_myc1_*tetO*-associated UTR or leaderless transcripts. The promoter sequence upstream of the transcription start site (TSS) was identical for all constructs; this suggests that sequences downstream of the TSS contribute to the *sigA* promoter, or otherwise positively influence production of the *sigA* transcript. This result highlights the complexity of bacterial transcription and advises caution when using TSS data to predict promoters.

The relative efficiency of leaderless vs. leadered translation in mycobacteria has not been experimentally established. Proteomics data from *M. tuberculosis* suggested that proteins encoded on leaderless transcripts were less abundant than those encoded on leadered transcripts with evident SD sequences, but this difference appeared to be explained by differences in mRNA levels [43]. A subsequently-reported quantitative proteomics dataset [60] allows for a more rigorous assessment of the relationship between mRNA abundance and protein abundance in *M. tuberculosis*. When comparing leaderless genes to leadered genes with a single TSS, we found that leaderless genes indeed had on average slightly but significantly lower levels of both mRNA and protein (Mann-Whitney tests for both, *p* < 0.01). However, the relationships between mRNA abundance and protein abundance did not differ for these two groups, consistent with the idea that there is no global difference in translation efficiency for leadered vs. leaderless transcripts. The small number of controlled comparisons we report here support that idea; transcripts with the p_myc1_*tetO*-associated UTR were translated more efficiently than leaderless transcripts, but a transcript with the *sigA* UTR was translated with similar efficiency as its leaderless counterpart. Notably, the difference in translation efficiency between the two leadered transcripts might be attributable to differences in secondary structure rather than differences in favorability of the SD sequences.

A positive correlation between mRNA half-life and translation efficiency was reported for *E. coli* [62], consistent with the idea that translation may protect mRNAs from degradation. We did not observe such a correlation without our set of five transcripts, indicating that translation efficiency is not the primary driver of the variability in half-lives we observed. However, a broad analysis of this relationship in mycobacteria is warranted.

## METHODS

### Strains and culture conditions

*Mycobacterium smegmatis* strain *ΔMSMEG_2952* [63] and its derivatives (Table 1) were grown in Middlebrook 7H9 medium with albumin/dextrose/catalase supplementation (ADC; final concentrations, 5 g/L bovine serum albumin fraction V, 2 g/L dextrose, 0.85 g/L NaCl, and 3 mg/L catalase), 0.2% glycerol, and 0.05% Tween 80. Cultures were shaken at 200 rpm and 37°C to an optical density at 600 nm (OD_600_) of ∼0.8 at the time of harvest.

**TABLE 1.** Strains and plasmids used.

| Plasmid | Strain | Characteristics |
| --- | --- | --- |
| - | ΔMSMEG_2952<sup>1</sup> | *mc2155*, MSMEG2952::*hyg<sup>r</sup>* |
| pSS303 | SS-M_0486 | P<sub>myc1</sub>*tetO* promoter + P<sub>myc1</sub> 5' UTR + *yfp-6xHis* |
| pSS309 | SS-M_0489 | P<sub>myc1</sub>*tetO* promoter + *sigA* 5' UTR + first 54 nt of *sigA* + *yfp-6xHis* |
| pSS310 | SS-M_0493 | P<sub>myc1</sub>*tetO* promoter + no 5' UTR + *yfp-6xHis* |
| pSS314 | SS-M_0497 | P<sub>myc1</sub>*tetO* promoter with a deletion of nt -53 through -1 + *yfp-6xHis* |
| pSS316 | SS-M_0521 | P<sub>myc1</sub>*tetO* promoter + Δ1GTG *sigA* + *yfp-6xHis* |
| pSS335 | SS-M_0524 | P<sub>myc1</sub>*tetO* promoter + Δ2GTG *sigA* + *yfp-6xHis* |
| pSS359 | SS-M_0623 | P<sub>myc1</sub>*tetO* promoter + P<sub>myc1</sub> 5' UTR + first 54 nt of *sigA* + *yfp-6xHis* |
| pSS360 | SS-M_0626 | P<sub>myc1</sub>*tetO* promoter + no 5' UTR + first 54 nt of *sigA* + *yfp-6xHis* |
| pSS365 | SS-M_0629 | P<sub>myc1</sub>*tetO* promoter with a deletion of nt -53 through -1 + first 54 nt of *sigA* + *yfp-6xHis* |
| pSS384 | SS-M_0636 | P<sub>myc1</sub>*tetO* promoter with a deletion of nt -53 through -1 + *sigA* 5' UTR + *yfp-6xHis* |
| pSS385 | SS-M_0639 | P<sub>myc1</sub>*tetO* promoter with a deletion of nt -53 through -1 + P<sub>myc1</sub> 5' UTR + first 54 nt of *sigA* + *yfp-6xHis* |

<sup>1</sup> Strain source: [63]

**TABLE 2.**
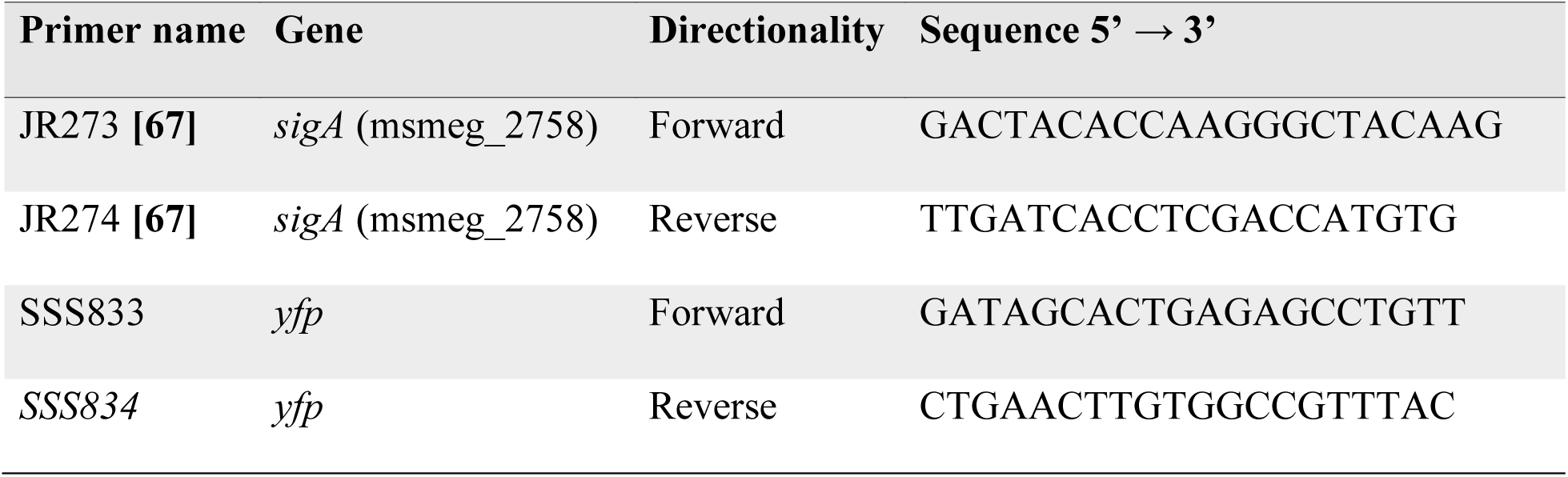
Primers for qPCR.

### Plasmid construction

Plasmid pSS303 was built on a backbone derived from pGH1000A [64] by inserting a *yfp* cassette containing the gene sequence of a YFP reporter (sfYFP, obtained from Ivy Fitzgerald and Benjamin Glick) with a 6x His tag at the C-terminus (complete amino acid sequence: MASDSTESLFTGVVPILVELDGDVNGHKFSVRGEGEGDATNGKLTLKLICTTGKLPVPWPTLVTTLGYGVQCFARYPDHMKQHDFFKSAMPEGYVQERTITFKDDGTYKTRAEVKFE GDTLVNRIELKGIDFKEDGNILGHKLEYNFNSHNVYITADKQKNGIKANFKIRHNVEDGGVQLADHYQQNTPIGDGPVLLPDNHYLSYQSKLSKDPNEKRDHMVLLEFVTAAGITHGSSGSSGCHHHHHH). Two synthetic transcriptional terminators were inserted flanking the cassette: *tsynA* [65] upstream and *tt*_*sbi*_*A/B* [66] downstream. Transcription was initiated by the P_myc1_*tetO* promoter, which was constitutively active in our strains due to the absence of the corresponding tet repressor [56]. All constructs (pSS303 and derivatives noted in Table 1) were built using NEBuilder HiFi DNA Assembly Master Mix (E2621). Each assembled plasmid was integrated in *M. smegmatis ΔMSMEG_2952* [63] at the Giles phage site, and selected with 200 µg/mL hygromycin.

### Cell fixation and flow cytometry

1.5 mL aliquots of *M. smegmatis* cultures were pelleted, resuspended in 500 µL 2% paraformaldehyde in PBS, and incubated at room temperature for 30 min. Cells were rinsed twice using 900 µL phosphate saline buffer (PBS) + 0.1% Tween 20 and resuspended to a calculated OD_600_ of 15. Prior to flow cytometry analysis, cells were filtered using an 18-gauge 5µm filter needle and diluted with Middlebrook 7H9 to an OD_600_ of 0.015. YFP fluorescence intensity was measured per manufacturer’s instructions using a BD Accuri C6 flow cytometer collecting 100,000 events per sample (Fig. 1B and C), or a BD LSR II flow cytometer collecting 50,000 events per sample (Fig. 2B, 3B and 4B), using appropriate controls and thresholds. FlowJo v10.6 was used to draw tight forward scatter and side scatter gates to limit analysis to similarly sized cells, and GraphPad Prism 8 was used for statistical analysis.

### RNA extraction and determination of mRNA abundance and stability

RNA extraction, measurement of mRNA abundance, and mRNA stability analysis from *M. smegmatis* cultures were conducted in biological triplicates as described in [15]. Briefly, mRNA abundance was measured by quantitative PCR (qPCR) using iTaq SYBR green (Bio-Rad) on an Applied Biosystems 7500 with 400 pg of cDNA and 0.25 µM each primer in 10 µL reactions. Cycle parameters were 95°C for 15 seconds, and 61°C for 60 seconds. Primers used to determine mRNA abundance were JR273 (5’ GACTACACCAAGGGCTACAAG 3’) and JR274 (5’ TTGATCACCTCGACCATGTG 3’) [67] for *sigA*; and SSS833 (5’ GATAGCACTGAGAGCCTGTT 3’) and SSS834 (5’ CTGAACTTGTGGCCGTTTAC 3’) for *yfp*.

For mRNA stability analysis, 5 mL *M. smegmatis* cultures were treated with rifampicin at a final concentration of 150 µg/mL to halt transcription, and snap-frozen in liquid nitrogen after 0, 1, 2, or 4 min. Abundance over time was determined for *sigA* and *yfp* using qPCR and used to estimate mRNA half-lives as described above and in [15]. As is typical for bacteria, plotting log2 abundance over time produced a biphasic decay curve consistent with a period of faster exponential decay, followed by a period of much slower exponential decay. Because the initial phase is more likely to reflect decay rates in the cell prior to perturbation of cellular physiology with rifampicin, only the initial, steeper slope was used for mRNA half-life calculations (0, 1 and 2 min for strain SS-M_0489; and 0 and 1 min for strains SS-M_0493 and SS-M_0626).

### Calculation of transcription rates

The rate of transcription was estimated as described in [57]. Briefly, transcription rate (*V*_*t*_) is described as:

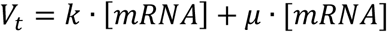

Where [mRNA] is a given transcript’s concentration, *µ* is the growth rate of the cells, and *k* is the degradation rate constant.

### Protein extraction and BCA assay

*M. smegmatis* cells were pelleted, rinsed three times with Middlebrook 7H9, 0.2% glycerol and 0.05% Tween 80 at 4°C, resuspended in PBS + 2% SDS + protease inhibitor cocktail (VWR, #97063-972), and transferred to 2 mL disruption tubes (OPS Diagnostics; 100-µm zirconium lysing matrix, molecular grade). Cultures were lysed using a FastPrep-24 5G instrument (MP Biomedical) using four cycles of 6.5 m/s for 30 s, with 1 minute on ice between cycles. Samples were clarified by centrifugation at 21,130 × g at 4°C for 10 minutes, and the supernatant containing protein was recovered and stored at −20°C. Protein concentrations were calculated using the Pierce™ BCA Protein Assay (Thermo Scientific, #23225) according to the manufacturer’s instructions.

### Western blotting

Protein was normalized to the indicated masses in a final volume of 9 µL, combined with 4 µL of 4X Protein Loading Dye (200 mM Tris-HCl pH 6.8, 400 mM DTT, 8% SDS, 0.4% bromophenol blue, 40% glycerol) and heated to 95°C for 10 minutes. Using gradient gels (4–15% Mini-PROTEAN® TGX™ Precast Protein Gels,Bio-Rad, #4561086), the samples were electrophoresed for 60 min at 140 V and then transferred to a PVDF membrane. The membrane was incubated in Blocking Solution (PBS + 5% non-fat milk) for 30 minutes, and washed once for 5 min using Washing Buffer (PBS 1X buffer + 0.1% Tween 20). The membrane was probed with 1 µg/mL His-tag Antibody, pAb, Rabbit (Genscript, #A00174) in Blocking Solution for 60 min at room temperature. The membrane was then rinsed twice with Wash Buffer and once with 1X PBS, and incubated with Anti-Rabbit IgG–Peroxidase (Sigma-Aldrich, #A4914), 1: 30,000 in Blocking Solution, for 60 minutes at room temperature. The membrane was rinsed as previously described, and incubated with HRP substrate (Radiance Q, Azure Biosystems, #AC2101) as recommended by the manufacturer. Imaging was done using an Azure C200 imaging system (Azure Biosystems).

### Software

GraphPad Prism was used for all linear regressions and comparisons (GraphPad Software, La Jolla California USA, www.graphpad.com). The Srna program within Sfold was used for RNA secondary structure predictions [58, 59].

## AUTHOR CONTRIBUTIONS

TGN, LAR, and SSS conceived and designed the experiments. TGN and DVB performed the experiments. TGN, DVB, and SSS analyzed the data. TGN, DVB, and SSS wrote the manuscript.

## ACKNOWLEDGEMENTS

This work was supported by NSF CAREER award 1652756 to SSS. Thanks to Ivy Fitzgerald and Benjamin Glick for providing the sfYFP construct. We thank all members of the Shell lab for technical assistance and helpful discussions.

## SUPPLEMENTAL TABLE

**Supplemental Table 1**. *M. smegmatis* and *M. tuberculosis* 5’ UTRs between 6 and 300 nt in length.

**Figure S1.**
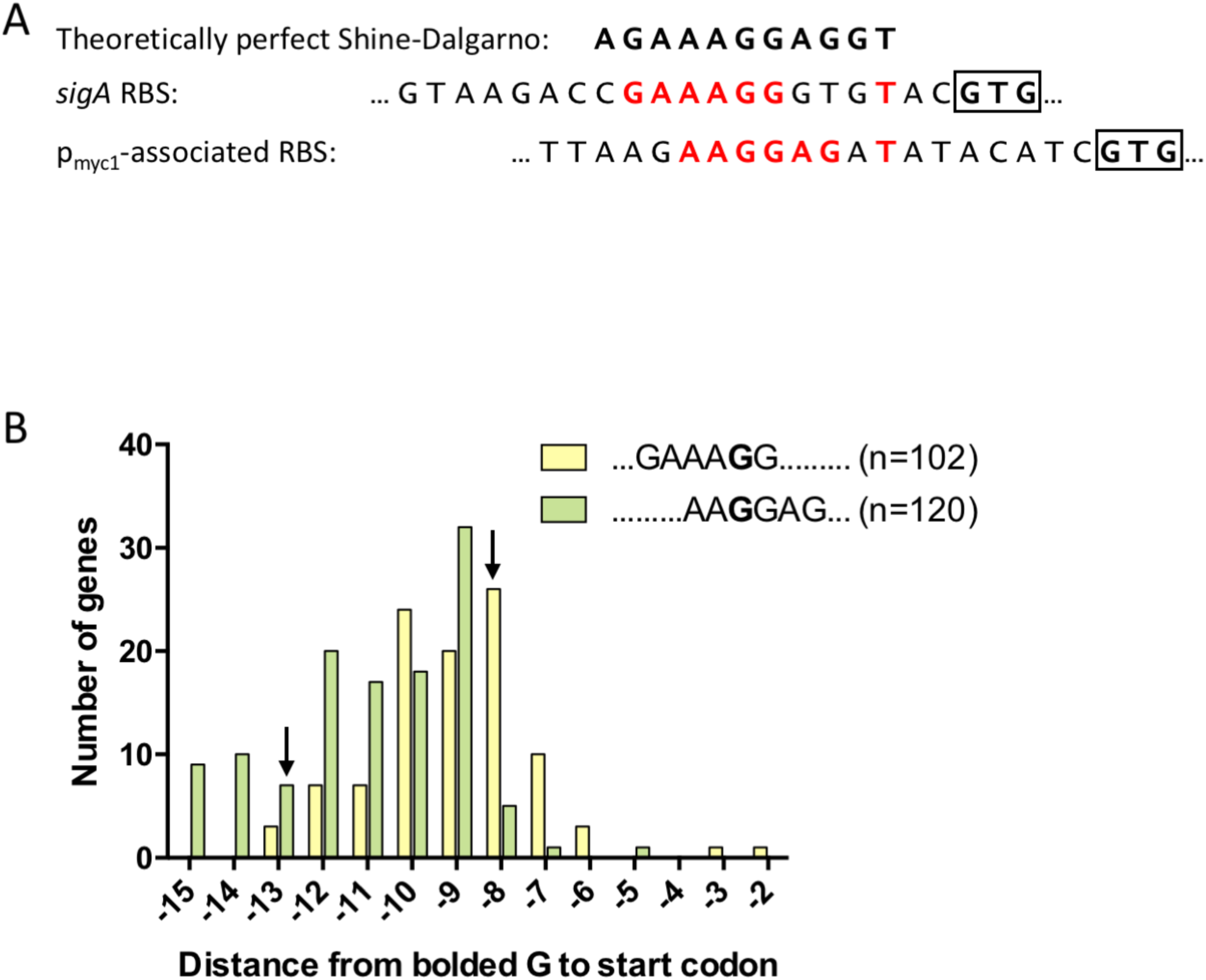
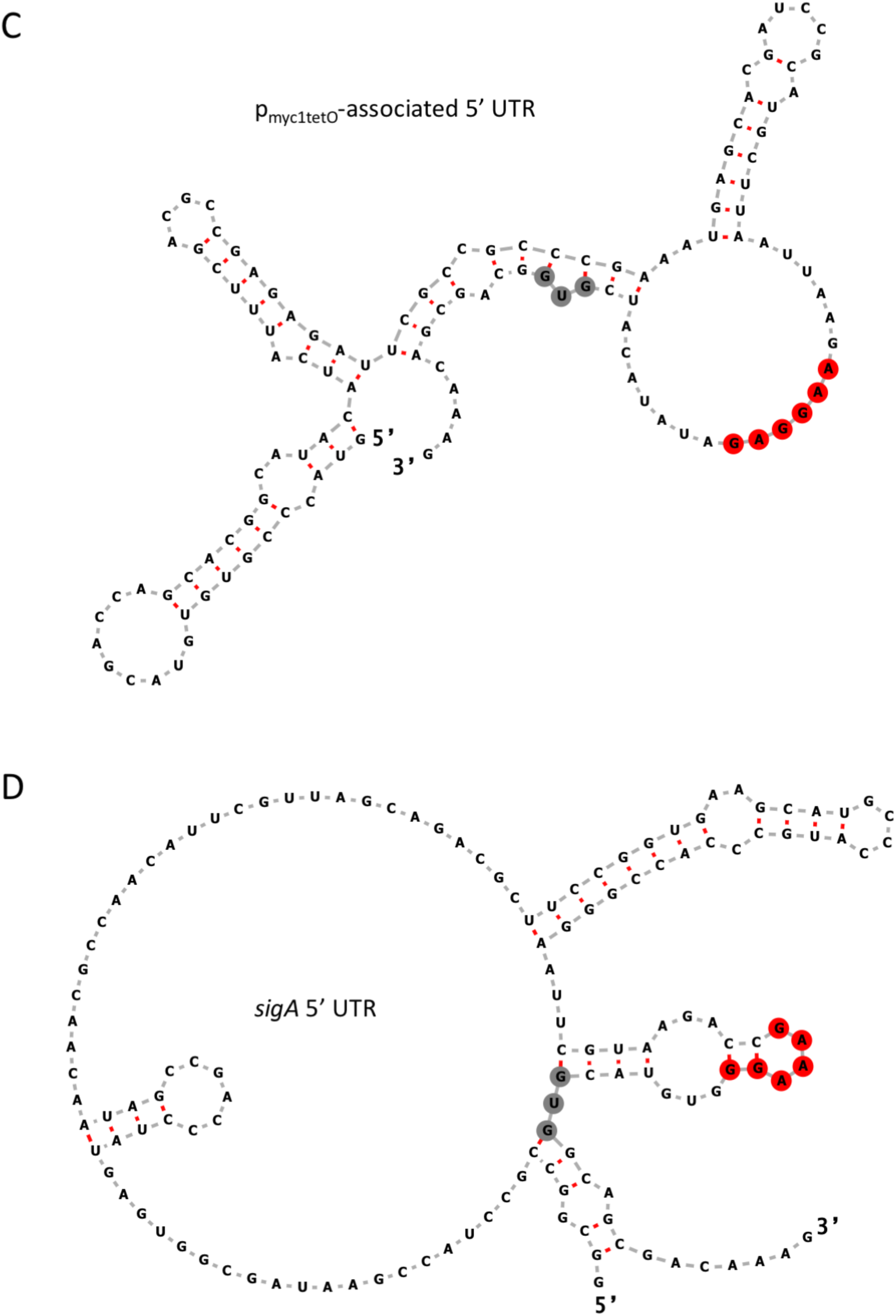
Comparison of Shine-Dalgarno (SD) sequences and predicted secondary structures for the *sigA* 5’ UTR and the p_myc1tetO_-associated 5’ UTR. **A:** The *sigA* and p_myc1_*tetO*-associated RBSs are shown aligned to the reverse complement of the 3’ end of the *M. smegmatis* 16S rRNA. Positions that match this theoretically perfect SD sequence are highlighted in red. Start codons are bolded and boxed. **B:** Distributions of SD-start codon spacings for all genes that have the indicated SD sequences in the *M. smegmatis* genome. Yellow indicates the *sigA* SD sequence and green indicated the p_myc1_*tetO*-associated SD sequence. Black arrows indicate the SD-start codon spacings for the *sigA* and p_myc1_*tetO*-associated SD sequences. **C-D:** Ensemble centroid predictions (Sfold, [59]) for secondary structures formed by the p_myc1tetO_-associated (C) and *sigA* (D) 5’ UTRs plus the first 15 nt of the *sigA* coding sequence. The predicted core SD sequences are highlighted in red. Start codons are highlighted in gray. The structures of the RBS regions were predicted to be the same when folding was performed using only the UTRs and start codons or using the UTRs and 54 nt of the *sigA* coding sequence.

